# Gene expression changes of seven stonefly species in responses to a latitudinal-environmental gradient

**DOI:** 10.1101/2020.09.22.309179

**Authors:** Maribet Gamboa, Yusuke Gotoh, Arnelyn D. Doloiras-Laraño, Kozo Watanabe

## Abstract

Latitudinal variation has been known to create strong selection pressure for genomic variation that enables the adaptation and survival of organisms. By altering gene expression patterns, organisms can modify their adaptive potential to heterogeneous environmental conditions along a latitudinal gradient; however, there is a gap in our understanding of how physiological consequences in wild species are affected and how changing environmental conditions act on multiple species. Here, we investigated how seven stream stonefly species sampled from four geographical regions in Japan differ in their responses to latitudinal variations by measuring gene expression (RNA-sequencing) differences within species and gene co-expression among species. We found that a large number of genes (622) were differentially expressed along the latitudinal gradient. The high species-specific gene expression diversity found at higher latitude regions was probably associated with low temperatures and high water discharge, which suggests the adaptive potential of stonefly specie. In contrast, similar gene expression patterns among species was observed at lower latitudes, which suggests that strong environmental stress occurs in warmer regions. Weighted gene co-expression network analysis (WGCNA) identified 22 genes with similar expression patterns among species along the latitudinal gradient. Among the four geographical regions, high differential expression patterns in the co-expressed genes from two regions were found, suggesting that the local environment strongly affects gene expression patterns among species in these regions. Respiration, metabolism, and developmental co-expressed genes exhibited a latitudinal cline, showing clear evidence of divergent adaptive responses to latitude. Our findings demonstrate that stonefly species are differentially adapted to local environmental conditions, and imply that adaptation in gene expression could be shared by multiple species under environmental stress conditions. This study highlights the importance of considering multiple species when evaluating the consequences of environmental changes on aquatic insect communities, and possible mechanisms to cope with environmental changes.

## 1. Introduction

In the face of climate change, a major research objective of conservation biology has become the determination of which species can adapt beyond the current physiological limitations (Hoffmann & Sgro, 2011; Bellard et al., 2012). Physiological change can improve an organism’s ability to cope with environmental changes and can be the basis for achieving evolutionary adaptation and persistence (Bijlsma & Loeschcke, 2005). Regardless of whether physiological plasticity or local adaptation is responsible for these changes, an understanding of the underlying molecular mechanisms involved in these changes are key to better predict how species will respond to climate change (Hoffmann & Sgro, 2011).

Genetic methods developed over the last decades have generated multiple approaches to monitor gene expression variations regarding the link between physiological and environmental changes (Evans & Hofmann, 2012). Variations in gene expression play an important role in the adaptive processes of populations (Oleksiak et al., 2002; Whitehead & Crawford, 2006), are highly heritable (Whitehead & Crawford, 2006), and can lead to speciation that reflects species-specific physiological requirements (Uebbing et al., 2016). Differentially expressed gene (DEG) analysis has been used to study gene expression extensively. DEG analysis uses statistical methods to evaluate normalized read count data, which reveals quantitative changes that have occurred in gene expression (Costa-Silva et al., 2017). Among several environmental factors examined in DEG studies (e.g., Teets et al., 2012; Kvist et al., 2013), the most remarkable gene expression changes associated with environmental change have been attributed to latitudinal gradients (Zhao et al., 2015; Juneja et al., 2016; Porcelli et al., 2016; Salazar et al., 2019).

Latitudinal gradients have been observed to influence adaptive gene expression divergence among populations (Holloway et al., 2007; Pavey et al. 2010; Manel et al., 2010) as latitudinal gradients is linked with environmental variations such as temperature, UV radiation, and humidity (e.g., Fraser, 2013). Some examples of differential expression patterns in latitude-related genes include the upregulation of the circadian clock and metabolic genes in rice (*Oryza sativa*) caused by temperature and solar radiation variations at higher latitudes (Nagano et al., 2012), the regulation of oxidative stress genes related to salinization changes in fish (*Platichthys flesus*) at increasing latitudes (Larsen et al., 2007), and the latitude-linked upregulation of immunity and reproductive-related genes in flies (*Drosophila* melanogaster, Juneja et al., 2016). Hence, latitudinal gradients are an ideal system to study the effects of environmental variation on DEG patterns of organisms. However, previous studies have focused on target species in laboratory-controlled experiments, but wild multiple species still need to be studied.

Comparative studies on gene expression across multiple species have revealed important insights into molecular pathways and gene synchronization between species (Arnone & Davidson, 1997). One of the challenges of multi-species analysis is identifying co-expressed genes, which are genes that show similar expression profiles among different species under the same trait or condition. Co-expression analysis can be used to discover sets of co-expressed genes found in multiple species, and to discover genes associated with a specific trait (Schadt et al., 2005). Weighted gene co-expression network analysis (WGCNA) is one widely utilized method (Stuart et al., 2003), and is built from gene expression data by calculating co-expression values in terms of gene pairwise similarity scores at a significant threshold. WGCNA can detect the genes associated with a particular trait or condition (Zhang & Horvath, 2005), and has been used to detect co-expressed genes in several taxa (e.g., eukaryotes, Martin & Fraser, 2018; mouse, Fuller et al., 2007; cattle, Keogh et al., 2019; plants, Wisecaver et al., 2017). These studies have revealed new insights into co-expression; however, WGCNA has been scarcely used to investigate whether co-expressed genes are associated with environmental factors, except for a few studies, such as those on fish (Bernal et al., 2020) and coral (Kenkel & Matz, 2016).

In contrast to the studies focused on DEG along the latitude-environmental gradient, the spatial distribution of gene expression diversity (i.e., the number of genes expressed within a species, Zhang et al., 2015) along the gradient has received less attention. Notably, changes in gene expression diversity can result from alterations in gene expression levels, the replacement of expressed genes by other genes, or the downregulation or upregulation of specific gene expressions across different habitats. Changes in gene expression diversity along a latitudinal gradient have been reported in microbial communities to be lower in polar than in non-polar waters because of possible ocean warming (Salazar et al., 2019), but they have not yet been studied for any other taxa.

In this study, we evaluated the spatial change of gene expressions in seven stonefly species along a latitudinal gradient in Japan. Stoneflies are aquatic insects considered to be more sensitive to environmental changes, such as low oxygen concentrations and high water temperatures than other aquatic insects (Prenda & Gallardo-Mayenco, 1999). Stoneflies are distributed worldwide, with higher species diversity at higher latitudes, and the species has a high diversity and ecological relevance in freshwater ecosystems around the world (DeWalt & Ower, 2019). The seven species selected (*Perlodini incertae, Haploperla japonica, Nemoura ovocercia, Taenionema japonicum, Stavsolus japonicus, Amphinemura longispina*, and *Eocapnia nivalis*) have been studied previously for their responses to a latitudinal gradient in Japan using genomic and proteomics approaches, which showed genome-wide signatures of adaptive divergence among populations along the latitude gradient (Gamboa & Watanabe, 2019) using double-digest restriction site-associated DNA sequencing, and a high oxygen-protein expression at higher latitudes using quantitative proteomics and protein differential expression analysis (Gamboa et al., 2017), but, specific gene functions and relative gene expression levels related to the gradient remain to be determined.

A recent global study on aquatic insects has suggested an increase in organism abundance over the years (Klink et al., 2020), but, human-mediated environmental changes and global warming threaten the survival of aquatic insect species (Gamboa, 2010; Sheldon, 2012). Understanding the adaptive potential of species (i.e., ability of species to respond to selection by means of molecular changes, Eizaguirre & Baltazar-Soares, 2014) might be assessed by variations of gene expression patterns along the environmental gradient, could be a robust and practical way to understand the mechanisms behind their fitness, prevalence, and survival. Studies on gene expression changes as a result of environmental stress in aquatic insects have been rarely conducted on selected genes related to thermal effects in the mayfly *Neocloeon triangulifer* (Chou et al., 2018), gene expression profiling to stream drying in the caddisfly *Micropterna lateralis* (Erzinger et al., 2019), and thermal adaptation in the cold stonefly *Lednia tumana*, *L*. *tetonica* and *Zapada* sp. (Hotaling, Shah et al., 2020) under controlled experiments, where other species have not yet been investigated.

Here, we focused on the adaptive mechanisms of wild stonefly species by detecting changes in the gene expression profiles of the hemolymph. The insect hemolymph plays a key role in insect immunity, embryo development, cytokines, antioxidant proteins, and oxygen protein transportation (Kanost et al., 1990; Burmester, 1999). For the seven species, patterns of gene expression were analyzed from samples collected over a latitudinal gradient across the Japan Archipelago. Specifically, we aimed to (1) determine whether the transcriptional responses within species were influenced by latitude, (2) assess the spatial distribution of gene expression diversity along the latitude gradient, and (3) identify common molecular mechanisms associated with the latitudinal adaptation response in different species. Our results highlight the utility of RNA-sequencing (RNA-seq) analyses to identify candidate genes that underline the among-family variations in survival required for the adaptive response of natural selection.

## 2. Methods

### 2.1. Sampling collection and processing

We collected seven stonefly species that occur commonly through four regions of Japan, to test the potential effects of the latitudinal gradient on gene expression patterns. Seven species of stream stoneflies (*P. incertae*, *H. japonica*, *N. ovocercia*, *T. japonicum*, *S. japonicus*, *A. longispina*, and *E. nivalis*) were selected to gather a broad multispecies perspective. These species were selected as they represent six taxonomical families that each have their own different biological requirements, such as feeding behavior and habitat preferences. Stonefly nymphs were sampled at 12 sampling sites across four geographical regions (Matsuyama, Gifu, Sendai, and Sapporo) with different climatic conditions. Samples were collected over a 2-week winter period, starting at the end of January 2015 (Table S1) using D-flame nets (mesh size = 250 μm). Specimens collected at the latest developmental stage were identified using the taxonomic key of Japanese aquatic insects (Kawai & Tanida, 2005). RNA and proteins were extracted simultaneously *in situ* by withdrawing the hemolymph from live individual specimens with a sterile syringe. The hemolymph was placed into TRIzol reagent (Ambion), and RNA was isolated according to the manufacturer’s instructions.

### 2.2. RNA-seq library preparation

The RNA extracted from TRizol was cleaned using the RNeasy MinElute kit (Qiagen). The quality and concentration of the RNA was checked using the 2100 Bioanalyzer (Agilent Technologies, Inc.) and Qubit flex Fluorometer (ThermoFisher), respectively. Sample pools were used for expression analyses to ensure sufficient high-quality RNA and to reduce variance in expression due to individual differences. Two biological replicates were included for each species at each geographical region. Each biological replicate was produced from the pools of an average of 18 sampled individuals, as suggested by Rajkumar et al. (2015). RNA-seq libraries were performed using the protocol proposed by Wang (2011), with a few modifications. Briefly, the total RNA was fragmented at 70 °C for 5-min (RNA fragmentation reagents; ThermoFisher), and then precipitated with 3 M sodium acetate, glycogen, and 100% ethanol at −20 °C for 60 min. The cDNA was synthesized using the SuperScript III kit (ThermoFisher) according to the manufacturer’s instructions and then purified with Ampure DNA beads (Beckman Coulter). The purified dsDNA product was end-repaired (New England Biolabs) and A-tailed (Klenow fragment, New England Biolabs) according to the manufacturer’s instructions. The resulting product was added to a mixture of 1 μl indexed adaptor, 6.75 μl ligase buffer, 0.25 μl T4 DNA ligase (New England Biolabs), and 2 μl of water at 27 °C for 15-min. The mixture was purified with Ampure DNA beads, mixed with uracil DNA glycosylase (Enzymatics), and incubated at 37 °C for 30-min. The library was amplified using a mixture of dsDNA-uracil product, 1 μl illumina paired-end primers (10 μM each), 5 μl Phusion High-Fidelity buffer, 0.25 μl dNTP (25 mM), 0.25 μl Phusion High-Fidelity DNA Polymerase, and 2.5 μl water. The PCR mix was incubated as follows: 98 °C for 30 s, 10 cycles of 98 °C for 10 s, 65 °C for 30 s and 72 °C for 30 s, and the final elongation at 72 °C for 5-min. The final library was purified with Ampure DNA beads. Sixty-four libraries with different indexes were normalized (approximately 10 ng per sample), pooled, and sequenced on one Hiseq 4000 lane of 100-bp paired-end reads at the Beijing Genomics Institute, China. One species (*Isoperla nipponica*) was discarded because of low raw read output data at three of the four geographical regions. Thus, seven species were used for downstream analysis.

### 2.3. De novo assembly and comparative analysis

Raw reads were filtered at a quality cut-off of 20 and then trimmed of adapter sequences using the FASTX-Toolkit (Gordon & Hannon, 2010). Reads trimmed to a size shorter than 50 bp were discarded. *De novo* transcriptome assembly was conducted on filtered reads for each sample using the Trinity assembler version 2.2.1 (Haas et al., 2013). Redundant and extremely low-expressed contigs (consensus regions from overlapping reads) were removed using the filter_fasta_by_resem_values.pl Trinity-utility. A separate *de novo* transcriptome assembly from the pooled biological replicates of all samples resulted in a lower mean contig; therefore, sample-specific assemblies were used for subsequent analyses.

Homologous genes (i.e., genes inherited from a common ancestor) within a species and orthologous genes (i.e., genes from different species descended from a common ancestor) among multiple species were established using two approaches. First, we used MCScan (Wang et al., 2012) to identify putative homologous regions by synteny relationships (the physical location of contigs on the same putative chromosome within a species) for all *de novo* assembly contigs from two biological replicates per species. The relationships were identified based on a pairwise gene comparison of BLAST multiple alignment (similarity > 60%) scores from the best hit. Genes with the best hits and shared synteny were defined as homologous, however, genes that were best hits, but not syntenic were also defined as homologous, because of the existence of possible genomic rearrangements, as suggested by Zhao et al. (2015). The orthologous search was performed by collecting all homolog-contigs from all species into a single matrix, wich was assembled using the merge mode by StringTie version 2.1.0 (Pertea et al., 2015). This approach used the BLAST multiple alignment file, thus, we converted the file to a BAM file using Blast2Bam (https://github.com/guyduche/Blast2Bam), and sorted using SAMtools (Li et al., 2009) as an input file for the assembly. We used the flags –b and –e. All the homologs and ortholog-contigs (hereafter named as genes) found were then used to compare gene expression patterns across samples and geographical regions.

### 2.4. Gene expression analysis and annotation

The read counts for all genes from each sample were quantified using the map-based mode of Salmon version 0.0.1 (Patro et al., 2017), and selecting the validateMapping option. The files derived from Salmon were processed with the edgeR version 3.10 package (Robinson et al., 2010) in R version 3.3 (R Core Team, 2014). Gene quantifications were normalized using the Trimmed of Mean of M-values (TMM) method using the calcNormFactors function, and the counts per million (CPM) reads mapped function based on both the p-value (< 0.05) and the log2 fold change (> 1). DEG analyses were performed independently for each of the seven species using the glm function. The Benjamin-Hochberg false discovery rate of 1% was applied using the p.adjust function. Two species, *A. longispina* and *E. nivalis*, failed to achieve high-expressed genes for the Matsuyama region and were, therefore, excluded from further analysis from this region only. Heatmaps were generated using the log2 average expression of genes by combining all species across four geographical regions with the heatmap R package.

A co-expression network analysis among genes between the samples was performed to identify genes among each species that were associated with latitude. WGCNA (Zhang & Horvath, 2005) was performed using the WGCNA R package (Langfelder & Hovath, 2008). We followed the tutorials for undirected WGCNA, which involves a Pearson’s correlation for all gene pairs across all samples, the construction of a similarity matrix of gene expression through a power function, and the hierarchical clustering of samples based on the correlation with latitude data. Threshold power tests for the WGCNAs were performed using power = 10; mergeCutHeight = 0.3; min ModuleSize = 30; and TOMType = signed. Latitude data were obtained based on the geographical distance between each pair of sampling sites using the Euclidean distance extracted from the geographical coordinates as proposed by Gamboa and Watanabe (2019) using the Vicenty Ellipsoid package (Karney, 2013) in R. The WGCNA identifies modules based on the hierarchical clustering of highly interconnected genes that are associated with latitude.

The putative function of all gene sets (DEG, WGCNA, and homolog species-specific) was matched against the National Center for Biotechnology Information non-redundant transcripts and protein database (BLASTx, evalue 1 e −3). A homology search was used to explore the whole database without a taxonomical filter first, and then a taxonomical filter was applied to the search result using arthropods, drosophila, and stonefly databases. We obtained four separate homology outputs and compared their functions. The protein-coding genes obtained were subsequently analyzed by Gene Ontology (GO) enrichment analysis. The homology search and GO were performed using Blast2go version 5.2.5 (Conesa et al., 2005). Transcript nucleotide sequences were reverse translated by an amino acid converter (https://www.bioinformatics.org/sms2/rev_trans.html), using the universal invertebrate codon code (https://www.kazusa.or.jp/codon/) to evaluate the false discovery rate by BLASTx search (https://blast.ncbi.nlm.nih.gov).

### 2.5. Proteomics data matching

Protein information for the seven species studied here was obtained from Gamboa et al. (2017). Proteins were used to find a match corresponding to the transcript-genes, to corroborate or improve gene function identification. The nucleotide sequences of the proteins were obtained through reverse translation with an amino acid converter (http://www.bioinformatics.org/sms2/rev_trans.htm) using the universal invertebrate codon code (http://www.kazusa.or.jp/codon/). The nucleotide sequences of the proteins were used to find the matching corresponding transcripts using LAST (Frith & Kawaguchi, 2015) with multiple alignments (MAFT; Katoh, 2013) and > 60 % similarity. The hierarchical classification of the putative gene functions obtained was integrated with DAVID Tools (Huang et al., 2009).

### 2.6. Latitudinal boundaries and drivers

The expression diversity per species and geographical region was determined to observe species-specific differences along the latitudinal gradient using gene expression (TMM-normalized CPM values) on Simpson’s diversity indices in the vegan R package (Oksanen et al., 2012), as proposed by Zhang et al. (2015). Analysis of variance (ANOVA) and Tukey’s honest significance difference (HSD) analyses were performed on pairwise comparisons of the diversity value per species in R. Additionally, the proportion of gene similarity between each pair of taxa was observed to determine shared genes along the latitudinal gradient by quantifying the proportional similarity, as proposed by Whittaker (1952).

Gene co-expression profile differences in the stonefly communities were further investigated by observing gene expression patterns between the geographical regions. Principal components analysis (PCA) was performed from the gene expression (TMM-normalized CPM values) of the co-expressed genes obtained by the WGCNAs using the vegan R package. A comparison of the functional responses of each geographical region was further investigated with heatmaps using a log2 average expression of co-expressed genes in the heatmap R package.

## 3. Results

Paired-end Illumina sequencing generated 117 million raw reads, with individual counts ranging from 16.7 to 25.3 million per sample (median = 19.9 million reads) in 56 samples (Table S2). A total of 5.7 −9.1 million reads per sample (median = 7.1 million reads) were mapped successfully for *de novo* assembly (Table S2), which generated 4506 contigs ranging from 534 to 1012 contigs per sample (Table S3). The homologous and orthologous gene search analyses identified 3078 genes (Table S3), including 101 species-specific homolog-contigs (Table S4). Following these analyses, the genes were quantified based on the reads counts retrieving 1736 of the 3078 genes (Table S3).

### 3.1. Gene expression differences along the latitudinal gradient

A total of 622 of 1736 genes were identified to be differentially expressed (p-value < 0.05, and log2 fold change >1; Table S5) across the four geographical regions. Among the seven species, *H*. *japonica, S. japonicus*, and *T. japonicum* displayed slightly higher numbers of DEGs (an average of 120 genes, and a range of 100-157 genes).

Gene expression changes along the latitudinal gradient were observed using heatmaps. The DEGs revealed that the largest differences in the gene expression profiles within a species where found in the different geographical regions (Fig. S1). All stonefly species showed high expression diversity at higher latitudes (ANOVA < 0.05, Tukey’s HSD < 0.05 per species; Fig. 1), suggesting that the dominant factor influencing expression diversity was the latitude gradient. We observed that at higher latitudes, each species displayed a larger number of species-specific genes and a lower gene similarity. These observations decreased with decreasing latitude (Fig. 2). At lower latitudes, the species exhibited high gene similarity between the species, low expression diversity, and low species-specific gene expression. This highlights the high similarity of the gene expression profiles between the species.

**Fig. 1.**
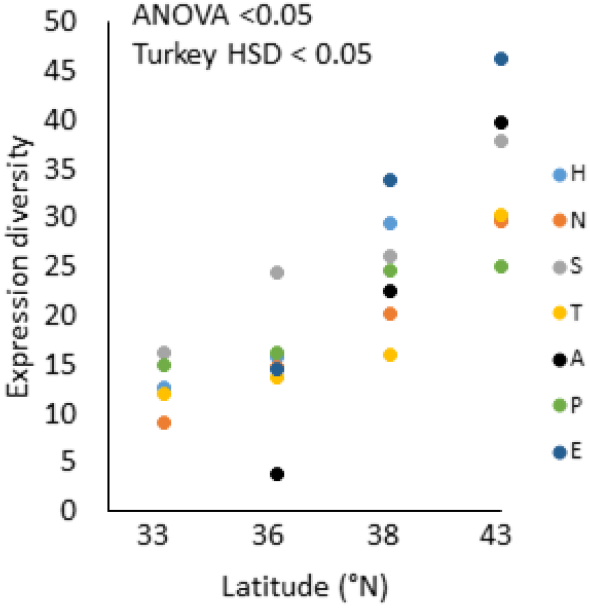
The expression diversity of stonefly species along the latitude gradient. H = *Haploperla japónica*, N = *Nemoura ovocercia*, S = *Stavsolus japonicus*, *T = Taenionema japonicum*, A = *Amphinemura longispina*, *P = Perlodini incertae*, E = *Eocapnia nivalis*.

**Fig. 2.**
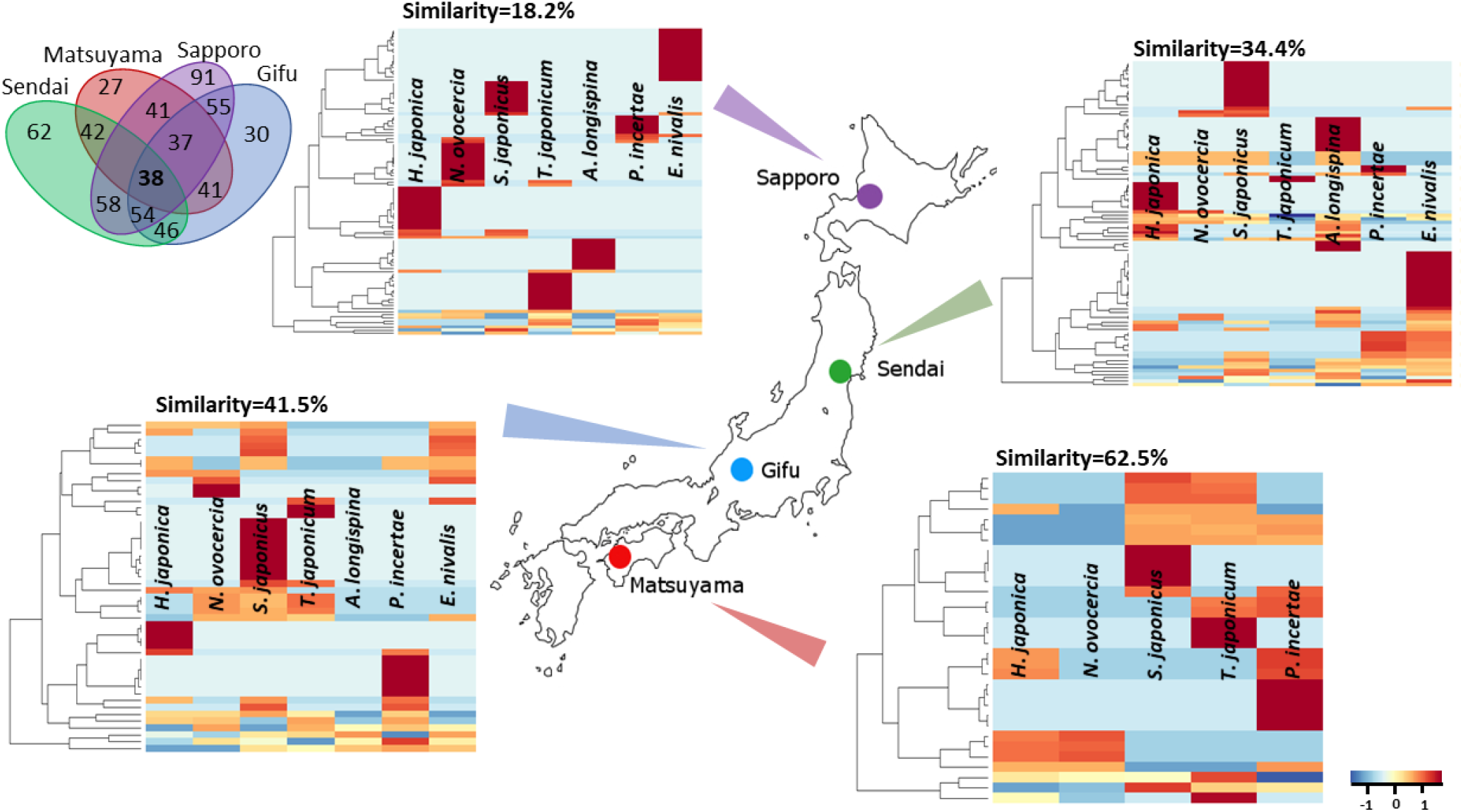
Differentially expressed genes (DEGs) of seven stonefly species across four geographical regions in Japan. Red colors represent highly expressed genes, while blue colors represent low expression (a color key is located on the lower right side). The proportion of gene similarity among species is shown at the top of the heatmap by each geographical region. The Venn diagram showing the number of DEGs found per and among geographical regions is located on the upper left side.

The BALSTx database was used to find the putative function of 723 genes (DEGs = 622, and homolog species-specific = 101). A blast was found for 174 genes, from which 89 genes were annotated to a function. We decided to retain five blasted genes without a matching gene ontology annotation for a possible function in the database. Among the 89 genes, 36% of the annotated genes were successfully matched to the proteomics data, and their associated proteins were obtained. The functions of several annotated genes that shared the same function were combined and a list of 30 total functions was created (Table S6). No annotations were found for species-specific or regional-specific genes, possibly because of the poor database for aquatic insect transcripts. The functions were divided into cellular components, molecular functions, and biological processes, and among these three, molecular functions were the most functionally diverse. Across the four geographical regions, the copper-binding functional gene related to the Hemocyanin protein was the most expressed gene for all species. The generation of precursor metabolites and energy functions had the highest number of genes with identical functions, suggesting that more than one related function is highly expressed between the four regions (Table S6).

### 3.2. Co-expression genes among stonefly species

The WGCNA highly correlated 22 genes with latitude, of the 3078 tested genes (Table S7). These 22 genes were found in 90% of the species in the four geographical regions and were among the expressed genes obtained through DEGs. PCA was performed to further investigate the gene expression patterns of these 22 genes between the stonefly species. The expression patterns clearly showed four clusters that represented the four geographical regions on the first two principal component axes (Fig. 3A). Approximately half of the total variance in the co-expressed gene among the samples was attributable to differences in latitudes (PCA, 59.6% of total variation). A single principal component axis separated three of the four geographical regions (Matsuyama, Sendai, and Sapporo). The largest species-gene expression variation within a geographical region was observed within Sapporo and Gifu.

**Fig. 3.**
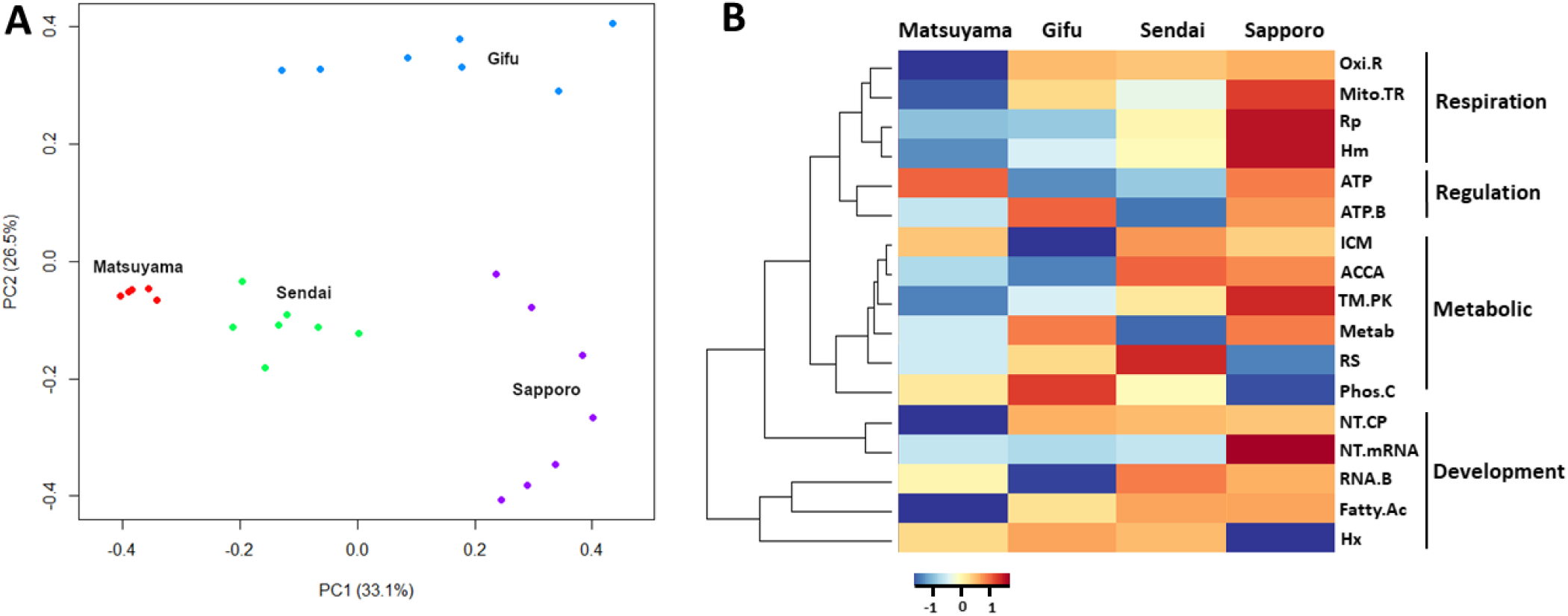
Comparison of geographical region co-expressed genes among seven species. (A) Principal component analyses of 22 co-expressed genes among four geographical regions. Red points represent Matsuyama, blue points Gifu, green points Sendai and purple points Sapporo. (B) Heatmap representing 17 of the 22 co-expressed genes that were functionally annotated by gene ontology (GO) analysis and blast. Oxi.R = oxidoreductase activity (GO:0016491), Mito.TR = mitochondrion, RP = respirasome (GO:0070469), Hm = copper ion binding (GO:0005507) Hemocyanin protein, ATP = ATP synthase (GO:0042773), ATP.B = ATP binding (GO:0005524), ICM = integral component of membrane (GO:0016021), ACCA = acyl carrier activity (GO:0000036), TM.PK = transmembrane receptor protein tyrosine kinase signaling pathway (GO:0007169), Metab = generation of precursor metabolites and energy (GO:0006091), RS = response to stimulus (GO:0042742), Phos.C = phosphatidylinositol phospholipase C activity (GO:0004435), NT.CP = nuclear-transcribed mRNA catabolic process (GO:0000956), NT.mRNA = nuclear-transcribed mRNA catabolic process, RNA.B = RNA binding (GO:0003723), Fatty.AC = fatty acid binding (GO:0005504), Hx = Hexamerin.

Among the 22 co-expressed genes, 17 genes were annotated to a function and associated with a protein (Table 1). Genes for respiration, regulation, metabolism, and development were obtained, and their expressions patterns differed between the four regions. Among these functions, respiration-related genes and some metabolic genes were upregulated at higher latitudes. In contrast, the development-related Hexamerin gene was upregulated at lower latitudes (Fig. 3B), which suggests that these genes display a latitudinal cline.

**Table 1.**
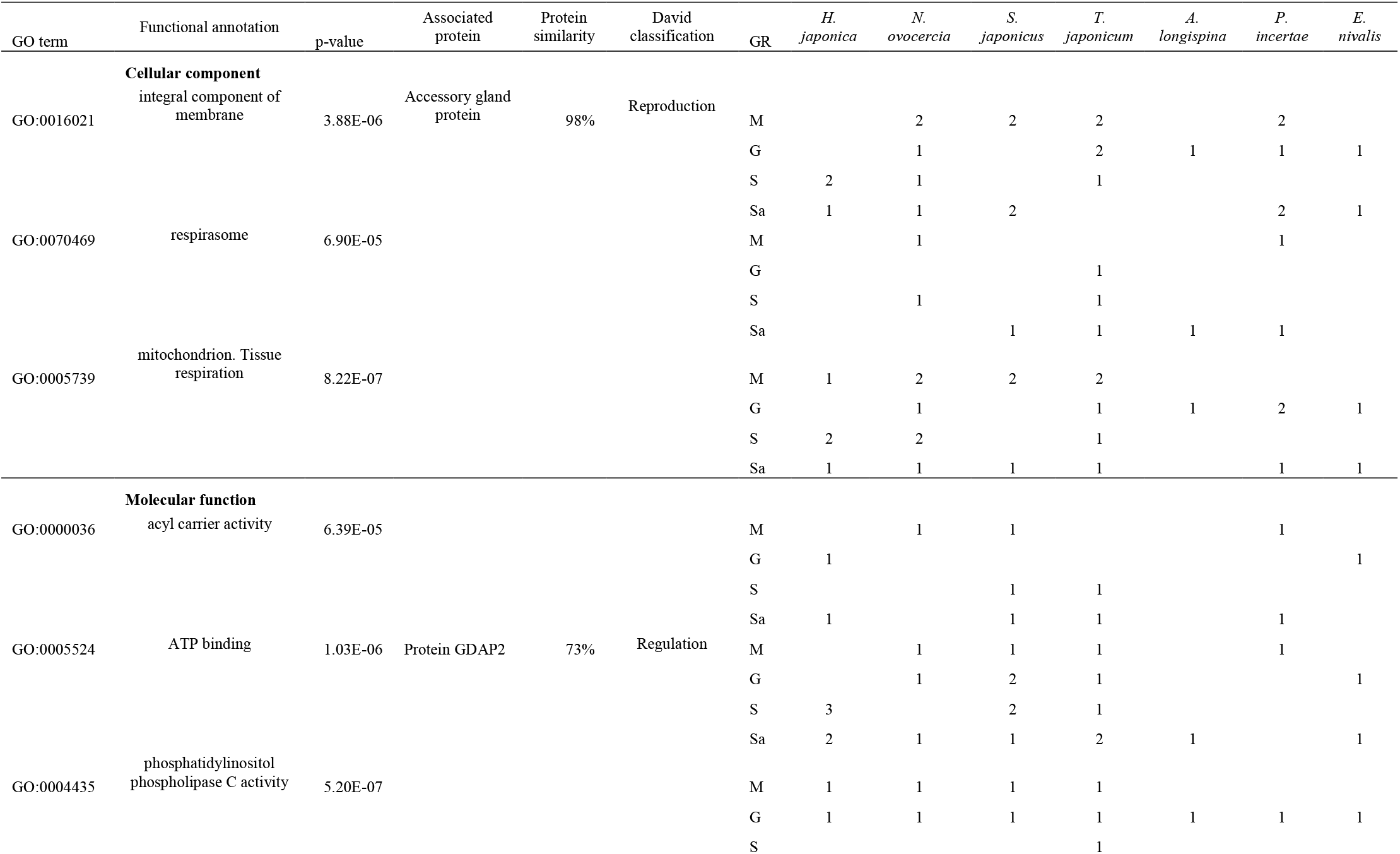
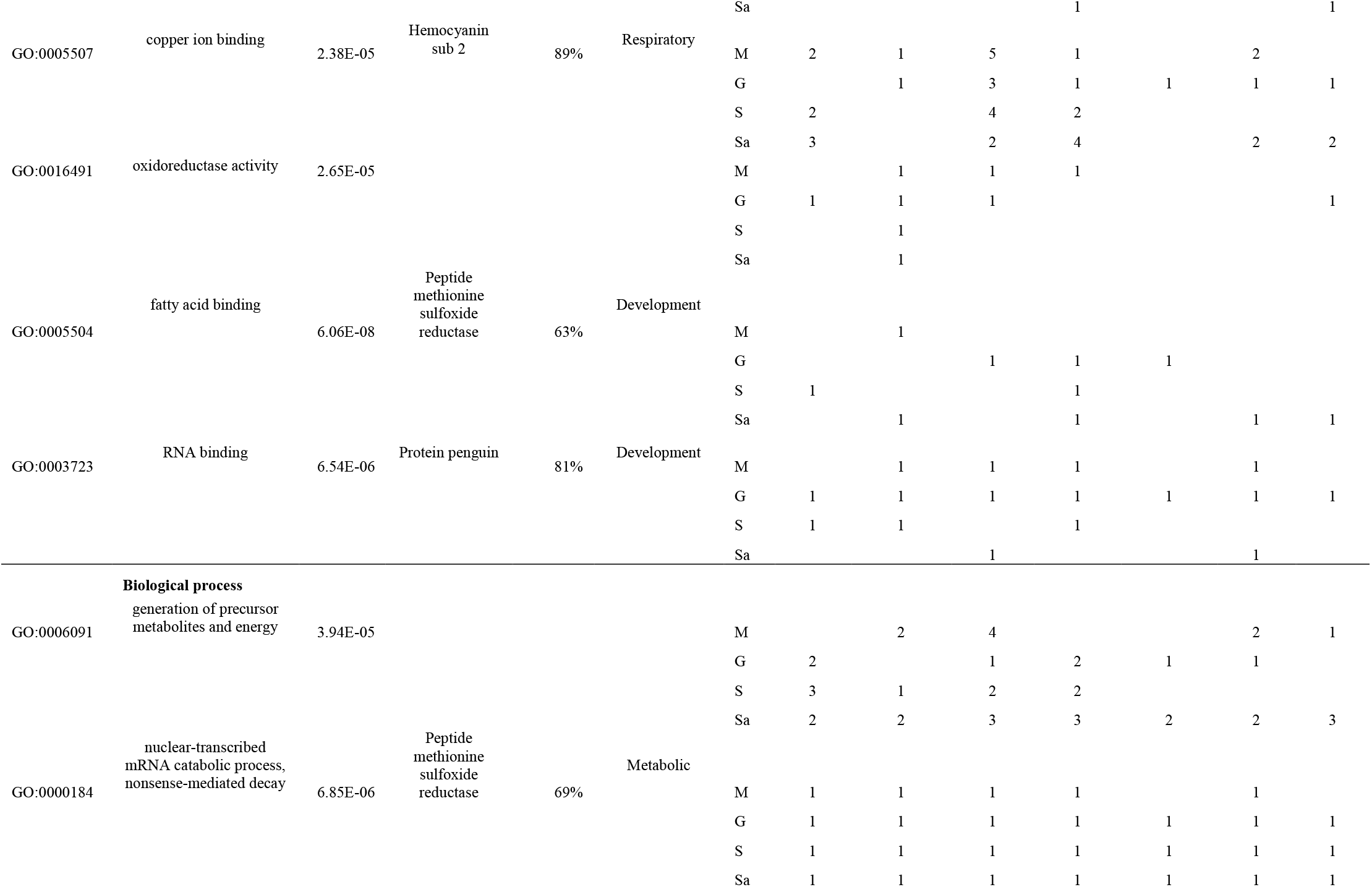
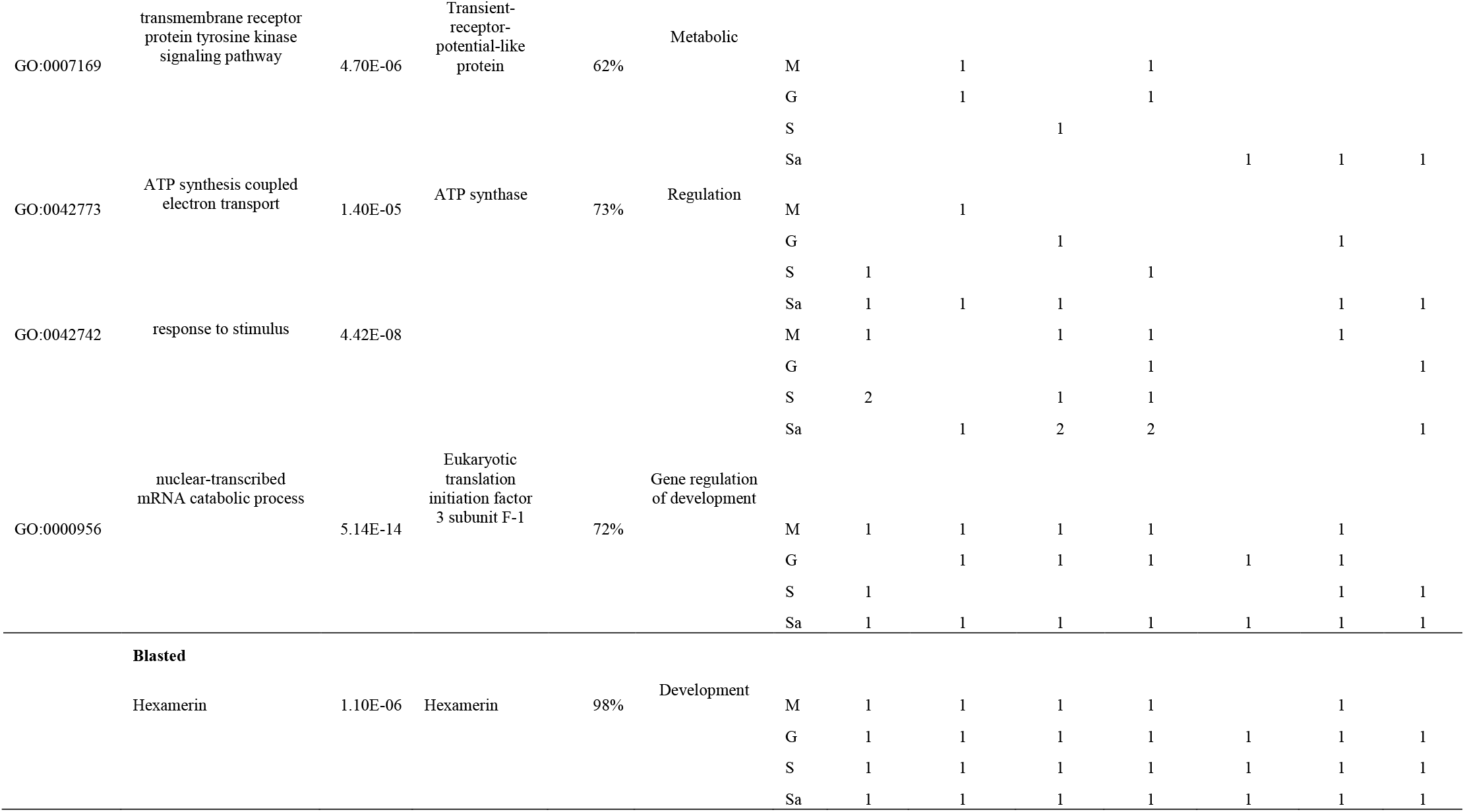
Functional annotated and blasted genes obtained from weighted gene co-expression network analysis (WGCNA) and their associated proteins. GO = Gene ontology, GR = geographical region, M = Matsuyama, G = Gifu, S = Sendai, Sa = Sapporo.

## 4. Discussion

Gene expression analysis is a powerful tool that can be used to investigate the physiological responses to environmental conditions or stressors, and can be indicative of environmental tolerance (Evans & Hofmann, 2012). Here, we explored RNA-seq to investigate the potential contribution of differential gene expression to latitudinal gradient adaptation across seven stonefly species. We observed that the latitudinal gradient influenced differential gene expression patterns in stonefly species and also enabled the co-expression of genes.

All seven stonefly species showed significant differential gene expressions along the latitudinal gradient, which concurs with previous studies on insects (*Drosophila* sp., Zhao et al., 2015; Juneja et al., 2016; Porcelli et al., 2016), plants (*Oryza sativa*, Nagano et al., 2012), and microbial communities (Salazar et al., 2019). Both local environmental conditions and the evolutionary history of the organisms could have played a significant role in this expression divergence. Environmental factors, such as temperature, rainfall, solar radiation, and humidity along the latitudinal gradient (Willig et al., 2003), are the main drivers of the selective pressures that result in varying gene expression profiles, which maximize fitness in the local environment (Hoffmann & Sgro, 2011). This was observed previously in the *D. melanogaster* adaptations that were driven by temperature and rainfall differences across latitudes (Zhao et al., 2015). Similarly, the evolutionary history of the organism could play a role in gene expression divergence, as observed previously in the marine snail, *Chlorostoma funebralis*. In this species, southern populations may employ heat-genes in anticipation to heat stress, an adaptation based on the evolutionary history of frequent heat exposure (Gleason & Burton, 2014). Stoneflies are generally weak fliers, with a limited airborne dispersal range within stream corridors, a long water stage during its immature stages (Stewart & Stark, 2008), and an evolutionary history linked with local environmental conditions (Gamboa et al., 2018; Gamboa & Watanabe, 2019). Thus, both local environmental conditions and evolutionary history could influence the stonefly gene expression differences we observed.

Differences in gene expression profiles among closely related species have been poorly documented; however, a common observation in gene expression studies indicates that the impacts of local environmental conditions depend on species-specific physiological traits (Oleksiak et al., 2002; Gleason & Burton, 2014; Somervuo et al., 2014; Huang et al., 2016; Kenkel & Matz, 2016; Bernal et al., 2020). We found that stonefly species at higher latitudes tended to increase their expression diversity and species-specific gene expression, but decrease their gene similarity to other species. This finding implies a stronger gene expression for species-specific physiological tolerance to higher-latitude regions. The northernmost region had the lowest atmospheric temperatures (annual mean 0.4 °C) and water temperatures (average 6.5 °C), but the highest water discharge (average 0.7 m^3^/s) (see Table S1). Although stoneflies are distributed worldwide, a high species diversity tends to occur at high latitudes (DeWalt & Ower, 2019), indicating that stoneflies can adapt to high-latitude environmental conditions. Likewise, a high water discharge can be a driving factor that increases stonefly species diversity, because the high disturbance levels that the habitats undergo allows for a high in-stream drift dispersal (Stewart & Stark, 2008). High species diversity is often linked with gene expression diversity, because high expression diversity promotes population persistence to the environment (Pavey et al., 2010). Moreover, high gene expression diversity at higher-latitude regions is probably associated with low temperatures and high water discharge, which could be an indicator of the possible adaptive potential of stonefly species.

Surprisingly, at lower latitude regions, five stonefly species (*P*. *incertae, H. japonica, N. ovocercia, T. japonicum*, and *S*. *japonicus*) displayed low gene expression diversity and high gene similarity, which could be an indication of environmental stress, as organisms tend to overlap their physiological responses to cope with stress. For example, the gene expression responses of five coral reef fishes to a heatwave showed an overlapping physiological response to high temperatures (Bernal et al., 2020). Similarly, the response of Atlantic salmon (*Salmo salar*) populations to thiamine deficiency (Harder et al., 2020), and the response of *Quercus lobata* oak populations to drought (Mead et al., 2019) showed physiologically overlapping responses at different sampling locations. All species studied here have different habitat preferences and feeding behaviors, therefore, we expected to see different gene expression profiles between the species, given their species-specific physiological requirements. Hence, the high interspecies similarity in gene expression profiles at lower latitudes could be interpreted as a strong signal of ongoing stress synchronization, which could be a coping mechanism to the environmental conditions.

In addition to the role of latitudinal-environmental gradients in gene expression differences within each species, we found 22 latitude-associated co-expressed genes among the seven species. To our knowledge, co-expressed genes among evolutionarily closed species in the context of a latitude gradient have not yet been studied, although co-expressed network analysis has been used to accurately correlate genes to environmental conditions, as was the case for five coral fish species to heat (Bernal et al., 2020), and as well coral (*Porites astreoides*) and their symbionts (*Symbiodlnlum* sp.) to local adaptation responses (Kenkel & Matz, 2016). The 22 co-expressed patterns clustered the stonefly communities clearly into four groups, which represented the four geographical regions. Among these regions, especially large differences in these 22 gene expression profiles were observed in the Gifu and Sapporo regions. The Gifu region has been observed previously as a hotspot of species diversity (Gamboa et al., 2018) and genetic diversity (Gamboa & Watanabe, 2019) for stonefly species, because of the geological formation history of the Japanese islands, while the Sapporo region has low water temperatures that are a suitable condition for high protein expression diversity among stonefly species (Gamboa et al., 2017). Moreover, gene expression may play an important role in the evolutionary process of adaptive divergence in stonefly species located in the Gifu and Sapporo regions based on their species-specific physiological requirements.

Among the 22 co-expressed genes, 17 genes were linked to a function and associated with a protein. The respiratory-related genes and some metabolic and developmental-related genes showed clear patterns of latitudinal cline in their expression patterns. The respiratory-related and metabolic genes were downregulated at decreasing latitudes. Respiratory-related genes are highly abundant in the hemolymph of stoneflies, especially Hemocyanin (Burmester, 2001). The Hemocyanin gene is highly expressed in normal oxygen conditions (Amore et al., 2009), but no expression was found in hypoxic (i.e., dissolved oxygen concentrations < 2 mg O_2_/l) environments (Gamboa, 2020), despite the high survival rate of stoneflies and their resistance to a lack of oxygen (Malison et al., 2020). This decreased expression of respiratory-related genes along the latitudinal gradient might indicate that stonefly species employ compensatory mechanisms to obtain oxygen when dealing with environmental stress (Gamboa, 2020), as observed previously in other insects (fly *D*. *melanogaster*, Gleixner et al., 2008; beetle *Tribolium castaneum*, Wang et al., 2018). Metabolic gene expression changes have been reported to be associated with temperature variations along a latitudinal gradient (Salazar et al., 2019). We identified metabolic genes related to the biosynthetic process, and cellular responses to a stimulus (acyl carrier activity and transmembrane receptor protein tyrosine kinase signaling pathway, see Table 1 and Fig. 3). Both genes have been observed to be associated with temperature fluctuations in other species, such as heat-moisture changes along a latitudinal gradient in the tree, *Populus trichocarpa* (Zhang et al., 2019). Thus, these variations in metabolic gene expression along the latitudinal gradient could be a clear signal of stonefly species adaptation.

In contrast, the expression of the developmental-gene, Hexamerins, was upregulated at lower latitudes. Hexamerins is a non-functional Hemocyanin involved in metamorphosis, molting, reproduction, and energy production, as it acts as a source of amino-acid storage (Burmester, 1999), and has been associated with the life-history changes and adaptation of individuals (Kvist et al., 2013), as well as faster development due to habitat fragmentation (Somervuo et al., 2014). High Hexamerins gene expression at high temperatures (Zhou et al., 2007) and low water oxygen concentrations (Gamboa, 2020) has been linked to phenotypic plasticity in insects. Therefore, the high expression of Hexamerins at lower latitudes could be associated with a developmental adaptation of stonefly species to certain environmental factors, such as high temperatures or low oxygen concentrations. Overall, our results suggest that co-expressed genes among the species studied lead to divergent adaptive responses to latitude, despite having the same gene function. Future studies that target aquatic insect communities, and the genes related to respiration, metabolism, and development that were found in this study could be used as monitoring genes that show sensitive responses to the changing environments associated with latitude.

Our study showed clear gene expression differences governed by latitudinal adaptation among stonefly species. Based on our results, we hypothesize that the impact of gene expression diversity on the gene expression profiles of stonefly species will be reduced more at a lower latitude than at higher-latitude regions, which could limit the adaptation of these species to warmer temperatures and be an indication of the potential consequences of climate change on their physiological tolerance. This hypothesis must be interpreted within the limitations of the data analyzed here, because it cannot account for the evolutionary adaptation of stonefly species to gradual changes over time. For example, among the seven species, predator and semi-predator species (*H*. *japonica, S. japonicus*, and *T*. *japonicum*) displayed a slightly higher number of DEGs than shredder species (*P. incertae, N. ovocercia, A. longispina*, and *E. nivalis*), without significant differences (t-test >0.05, data not shown). The differences in gene expression profiles between these two groups could be accentuated by changing seasons, during which behavior is affected greatly. Therefore, further studies need to resolve the long-term temporal dynamics of gene expression profiles among these stonefly species to improve our understanding of the contributions of gene expression changes within the context of environmental changes. Additionally, individual-based RNA-seq analysis rather than pooling samples could be used to better understand gene expression variations at the species level and could provide a different perspective of species adaptation. With the rapid advances in technology, new RNA extraction methods could also help improve the quality and quantity of samples for such studies. Similarly, increasing the number of samples and sampling locations in Japan is needed to further improve the interpretations of gene expression changes in stonefly species on a spatial scale.

## 5. Conclusions

Although latitudinal environmental variations has been studied often to better understand the local adaptation and environmental gradient impacts on species survival, our knowledge of freshwater ecosystem species remains scarce. The present study is the first to provide a spatial evaluation of the mechanisms underpinning community transcriptome changes in aquatic insects. We found that at lower latitudes, stonefly species tend to reduce gene expression diversity, probably as a method to cope with environmental stress. By contrast, at higher latitudes, the species displayed species-specific gene expression patterns that were probably linked with environmental tolerance and long-term evolutionary adaptation. Community co-expressed genes showed a latitudinal cline, wherein respiratory-related and metabolic genes cloud play an essential role in adaptation, and may be used for biomonitoring. Notably, our study could serve as a framework for future work on integrating temporary data to further investigate gene-ecosystem models that could improve ecosystem and climate policies.

## 6. Supplementary materials

The following supplementary materials are provided: supplementary tables 1–7, and supplementary figure 1.

## 7. Acknowledgments

For logistic support the fieldwork, we kindly thank Sueyoshi Masanao, Terutaka Mori, Ryota Kawanishi, and Kei Nukazawa.

## 10. Author contributions

M.G. and K.W. conceived the study; M.G. and K.W. acquired the funding; M.G. collected the field data; M.G. and Y.G. processed the data; M.G. and A.D.-L. analyzed the data; M.G. wrote the manuscript with insights from A.D.-L. and K.W.

## 11. Data availability statement

RNA-seq raw sequence reads are available from Genbank at the National Center for Biotechnology Information short-read archive database (BioProject accession no.: PRJNA647250)

## Supporting information

**Table S1.**
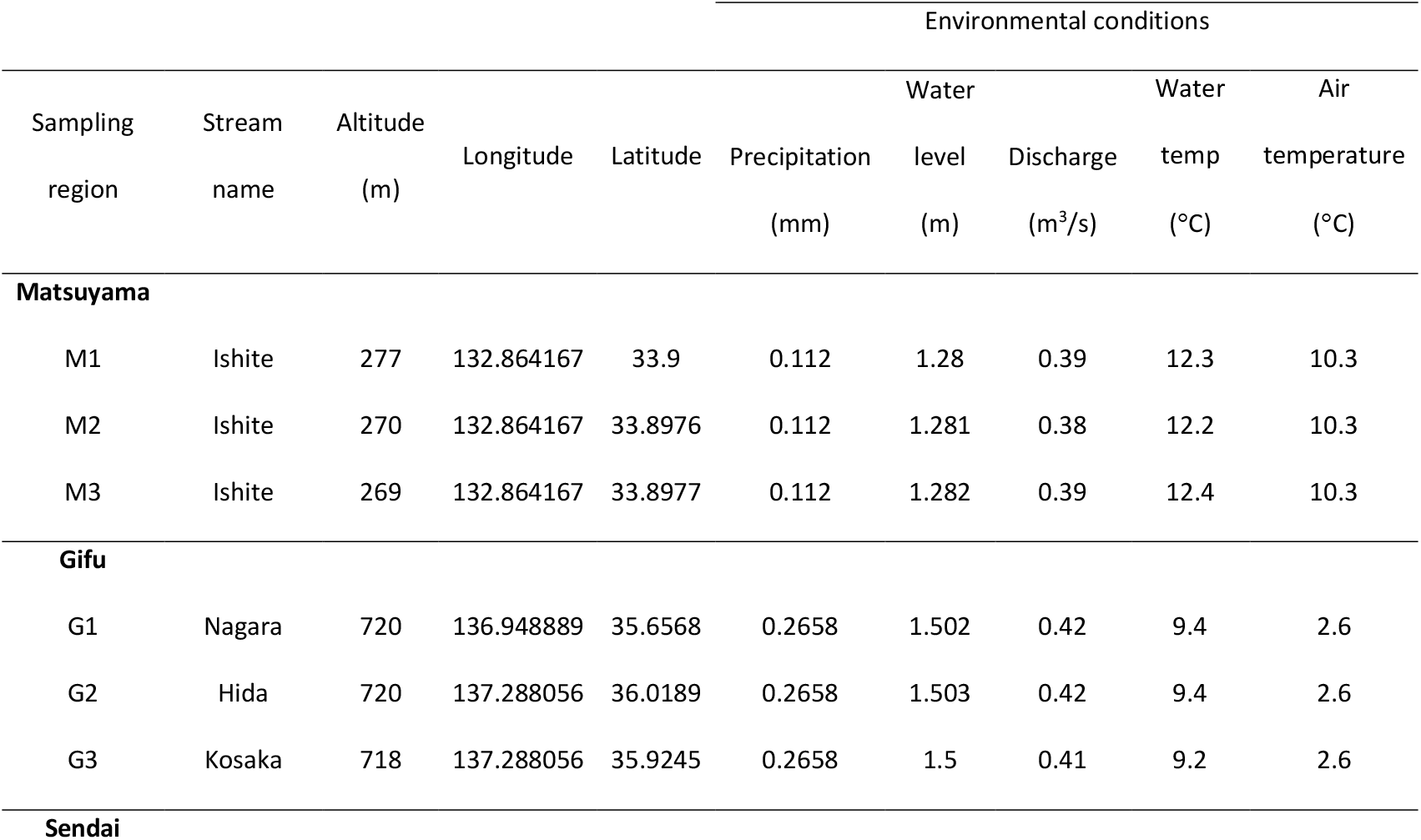
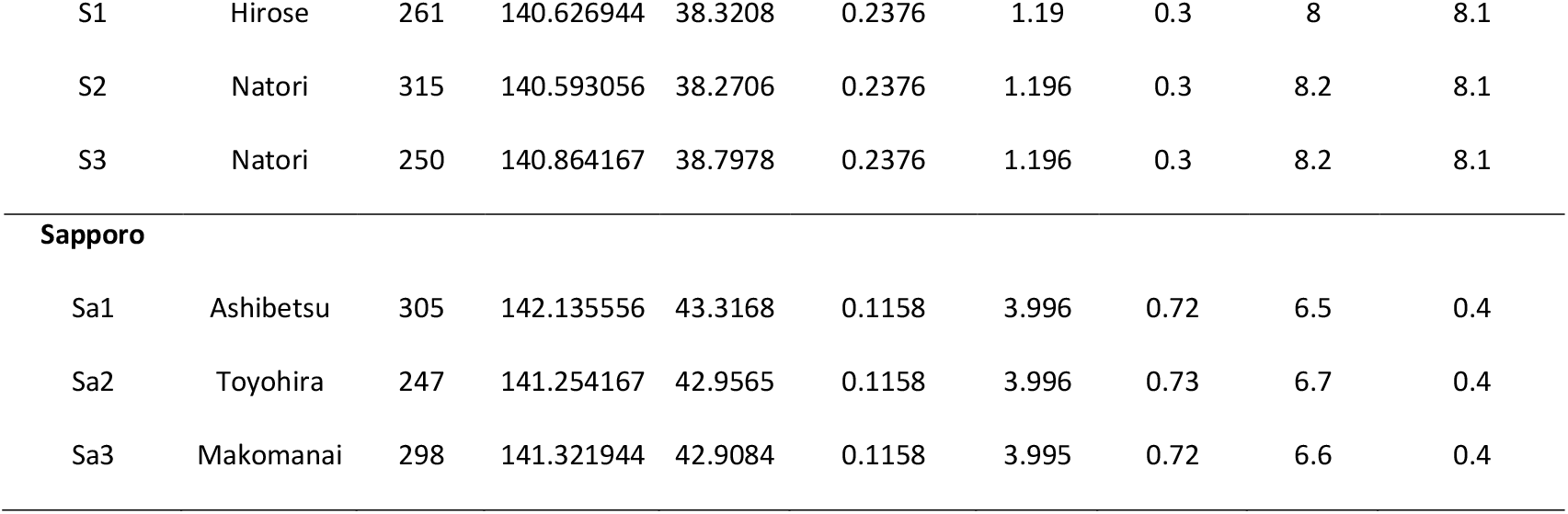
Sampling site information for the four geographical regions in Japan, including the average values for their environmental conditions.

**Table S2.** The number of total raw reads obtained following Illumina Hi-seq 4000 100bp paired-end sequencing and the number of reads mapped to *de novo* transcriptome assembly.

| Samples | Location | Raw reads | Mapped reads |
| --- | --- | --- | --- |
| *H. japonica* | Matsuyama.1 | 16,764,058 | 6,035,061 |
|  | Matsuyama.2 | 23,386,390 | 8,419,100 |
|  | Gifu.1 | 21,161,006 | 7,617,962 |
|  | Gifu.2 | 20,687,764 | 7,447,595 |
|  | Sendai.1 | 18,352,016 | 6,606,725 |
|  | Sendai.2 | 19,987,938 | 7,195,657 |
|  | Sapporo.1 | 19,642,970 | 7,071,469 |
|  | Sapporo.2 | 19,374,262 | 6,974,734 |
| *N. ovocercia* | Matsuyama.1 | 23,655,204 | 8,515,873 |
|  | Matsuyama.2 | 18,980,066 | 6,832,823 |
|  | Gifu.1 | 19,409,926 | 6,987,573 |
|  | Gifu.2 | 19,158,016 | 6,896,885 |
|  | Sendai.1 | 18,200,802 | 6,552,288 |
|  | Sendai.2 | 19,042,230 | 6,855,203 |
|  | Sapporo.1 | 17,888,558 | 6,439,881 |
|  | Sapporo.2 | 18,352,106 | 6,606,758 |
| *S. japonicus* | Matsuyama.1 | 18,244,020 | 6,567,847 |
|  | Matsuyama.2 | 20,829,064 | 7,498,463 |
|  | Gifu.1 | 24,091,696 | 8,673,011 |
|  | Gifu.2 | 17,266,168 | 6,215,820 |
|  | Sendai.1 | 17,363,874 | 6,250,994 |
|  | Sendai.2 | 17,504,798 | 6,301,727 |
|  | Sapporo.1 | 20,878,276 | 7,516,179 |
|  | Sapporo.2 | 20,941,176 | 7,538,823 |
| *T. japonicum* | Matsuyama.1 | 18,299,488 | 6,587,815 |
|  | Matsuyama.2 | 20,111,116 | 7,240,001 |
|  | Gifu.1 | 18,176,400 | 6,543,504 |
|  | Gifu.2 | 19,344,372 | 6,963,973 |
|  | Sendai.1 | 19,896,202 | 7,162,632 |
|  | Sendai.2 | 19,199,666 | 6,911,879 |
|  | Sapporo.1 | 18,730,870 | 6,743,113 |
|  | Sapporo.2 | 18,731,276 | 6,743,259 |
| *A. longispina* | Matsuyama.1 | 16,357,902 | 5,888,844 |
|  | Matsuyama.2 | 16,099,812 | 5,795,932 |
|  | Gifu.1 | 18,282,240 | 6,581,606 |
|  | Gifu.2 | 19,852,040 | 7,146,734 |
|  | Sendai.1 | 19,183,750 | 6,906,150 |
|  | Sendai.2 | 20,502,234 | 7,380,804 |
|  | Sapporo.1 | 19,291,826 | 6,945,057 |
|  | Sapporo.2 | 18,109,931 | 6,519,575 |
| *P. incertae* | Matsuyama.1 | 19,428,652 | 6,994,314 |
|  | Matsuyama.2 | 20,059,878 | 7,221,556 |
|  | Gifu.1 | 18,716,366 | 6,737,892 |
|  | Gifu.2 | 20,469,518 | 7,369,026 |
|  | Sendai.1 | 21,837,606 | 7,861,538 |
|  | Sendai.2 | 22,087,498 | 7,951,499 |
|  | Sapporo.1 | 21,996,856 | 7,918,868 |
|  | Sapporo.2 | 22,987,546 | 8,275,516 |
| *E. nivalis* | Matsuyama.1 | 21,144,935 | 7,612,176 |
|  | Matsuyama.2 | 20,332,101 | 7,319,556 |
|  | Gifu.1 | 21,077,804 | 7,588,009 |
|  | Gifu.2 | 22,796,724 | 8,206,820 |
|  | Sendai.1 | 21,941,086 | 7,898,790 |
|  | Sendai.2 | 25,000,668 | 9,000,240 |
|  | Sapporo.1 | 20,897,608 | 7,523,139 |
|  | Sapporo.2 | 25,300,918 | 9,108,330 |

**Table S3.** The number of genes obtained by performing *de novo* assembly, following homologs-orthologs search, and gene quantification analysis.

|  | *de novo* assembly | Homologs-Orthologs | Gene quantification |
| --- | --- | --- | --- |
| *H. japonica* | 560 | 358 | 209 |
| *N. ovocercia* | 618 | 459 | 292 |
| *S. japonicus* | 1012 | 820 | 451 |
| *T. japonicum* | 659 | 510 | 238 |
| *A. longispina* | 567 | 309 | 122 |
| *P. incertae* | 558 | 407 | 227 |
| *E. nivalis* | 532 | 215 | 197 |
| **Total** | **4506** | **3078** | **1736** |

**Table S4.** The number of species-specific genes obtained following homologous identification based on reciprocal BLAST analysis.

|  | Matsuyama | Gifu | Sendai | Sapporo |
| --- | --- | --- | --- | --- |
| *H. japonica* |  | 1 | 4 | 11 |
| *N. ovocercia* |  |  |  | 7 |
| *S. japonicus* |  | 4 | 11 | 6 |
| *T. japonicum* | 1 |  |  | 8 |
| *A. longispina* |  |  | 6 | 9 |
| *P. incertae* | 1 |  | 3 | 4 |
| *E. nivalis* |  |  | 12 | 13 |
| **Total** | **2** | **5** | **36** | **58** |

**Table S5.** The number of differentially expressed genes among stoneflies species (false discovery rate < 0.01).

|  | Matsuyama | Gifu | Sendai | Sapporo | Total DE genes |
| --- | --- | --- | --- | --- | --- |
| *H. japonica* | 9 | 46 | 14 | 31 | 100 |
| *N. ovocercia* | 14 | 11 | 32 | 18 | 75 |
| *S. japonicus* | 31 | 31 | 31 | 64 | 157 |
| *T. japonicum* | 19 | 26 | 23 | 35 | 103 |
| *A. longispina* |  | 10 | 12 | 18 | 40 |
| *P. incertae* | 23 | 11 | 26 | 24 | 84 |
| *E. nivalis* |  | 12 | 13 | 38 | 63 |
| **Total** | **96** | **147** | **151** | **228** | **622** |

**Table S6.**
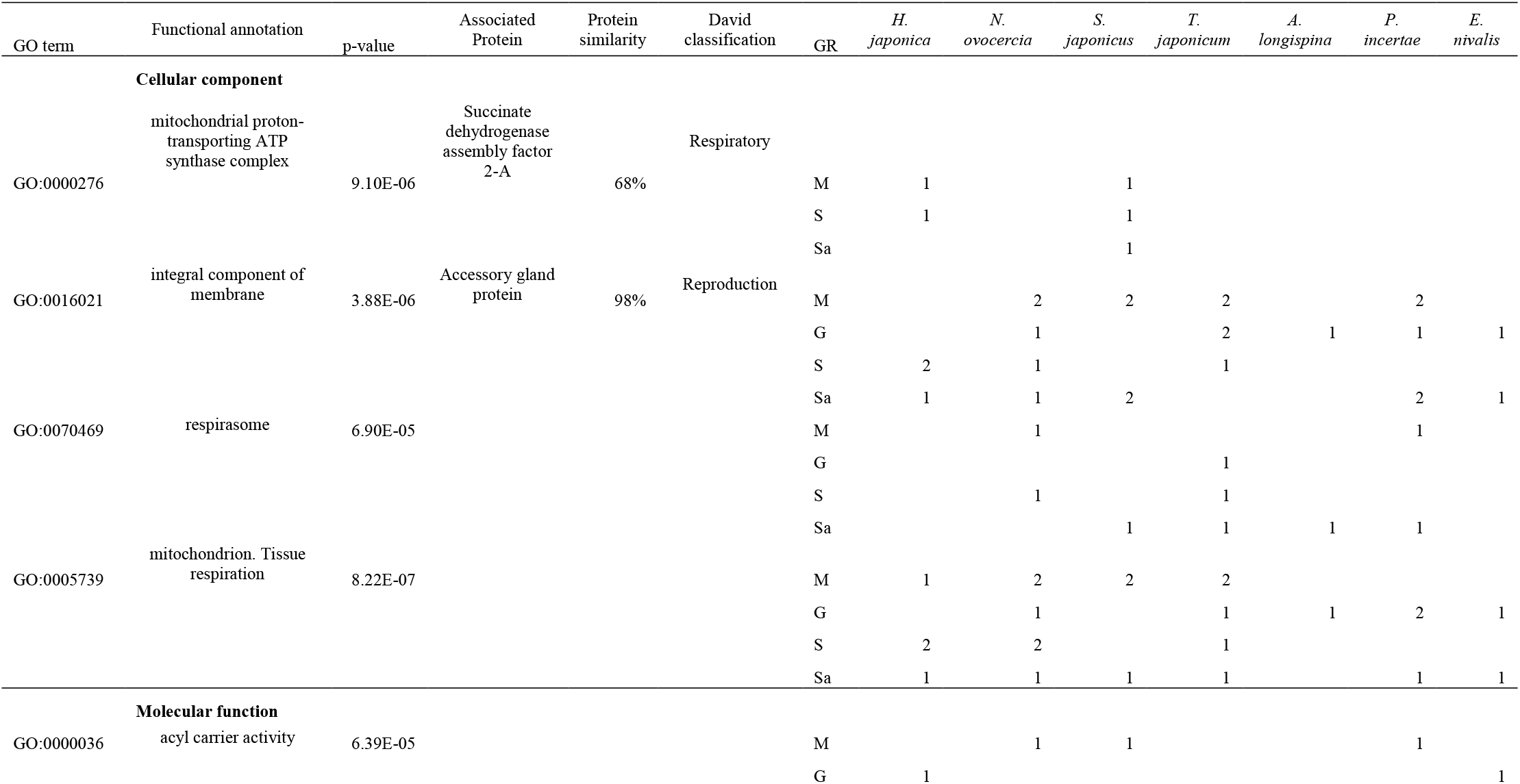
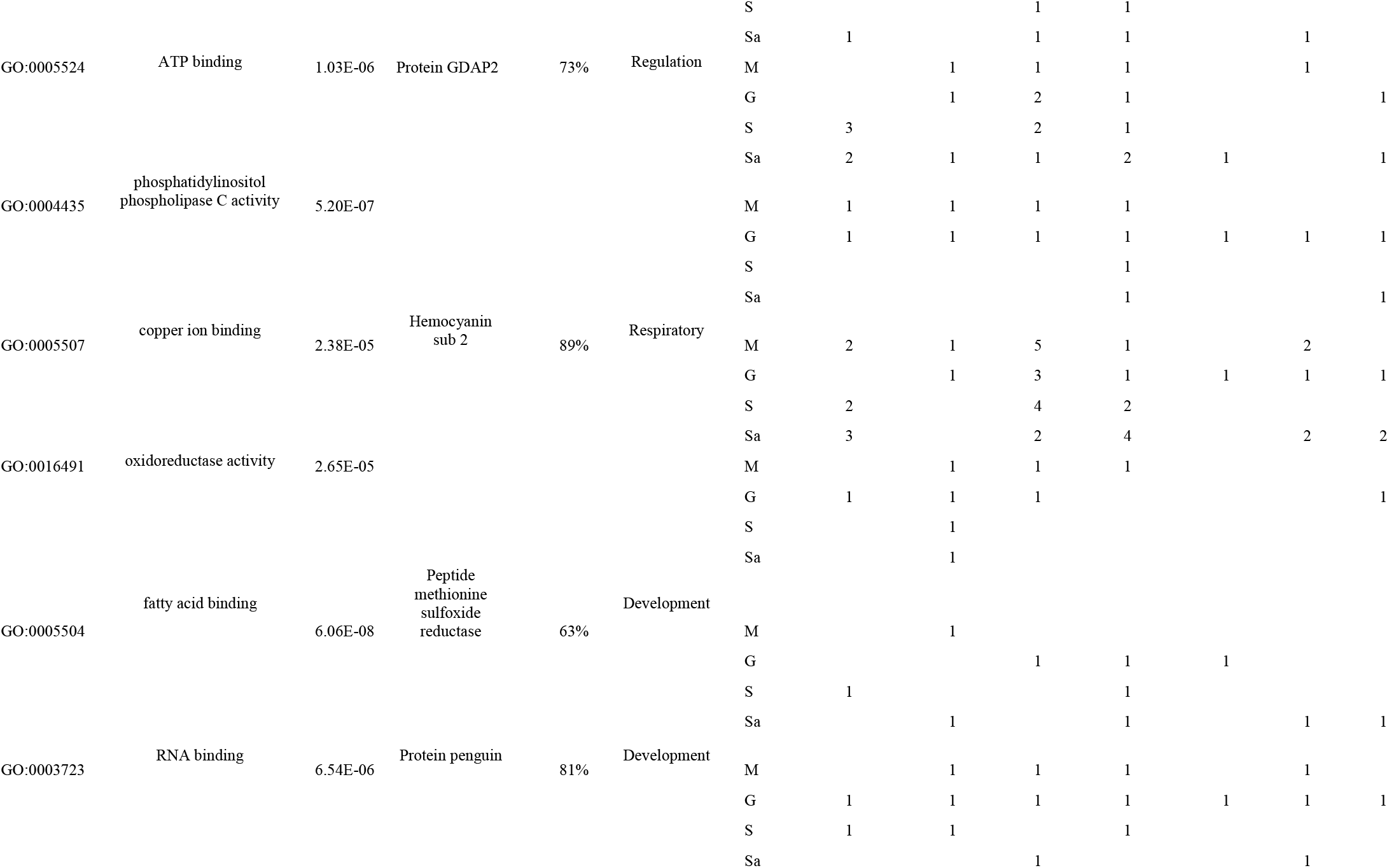
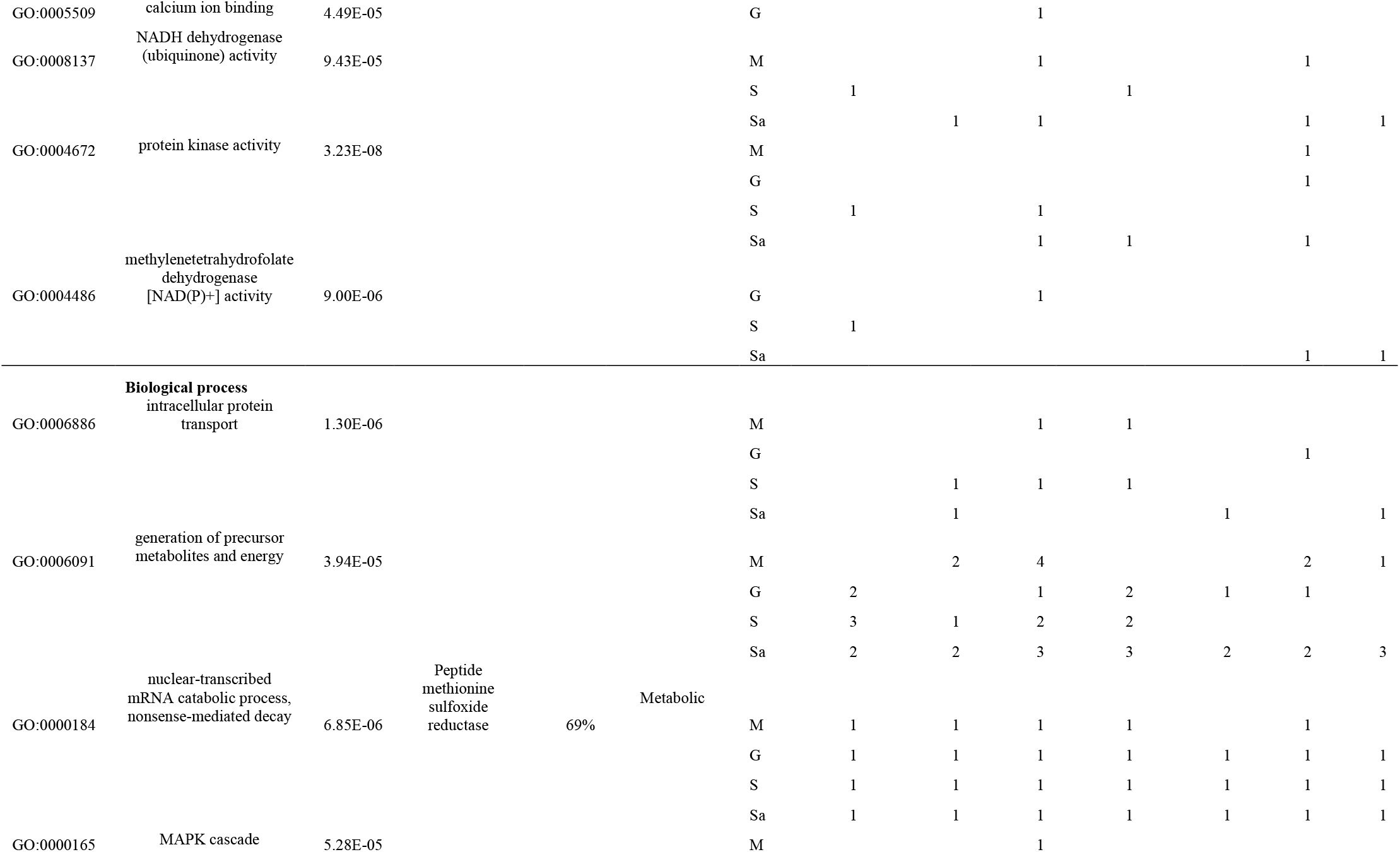
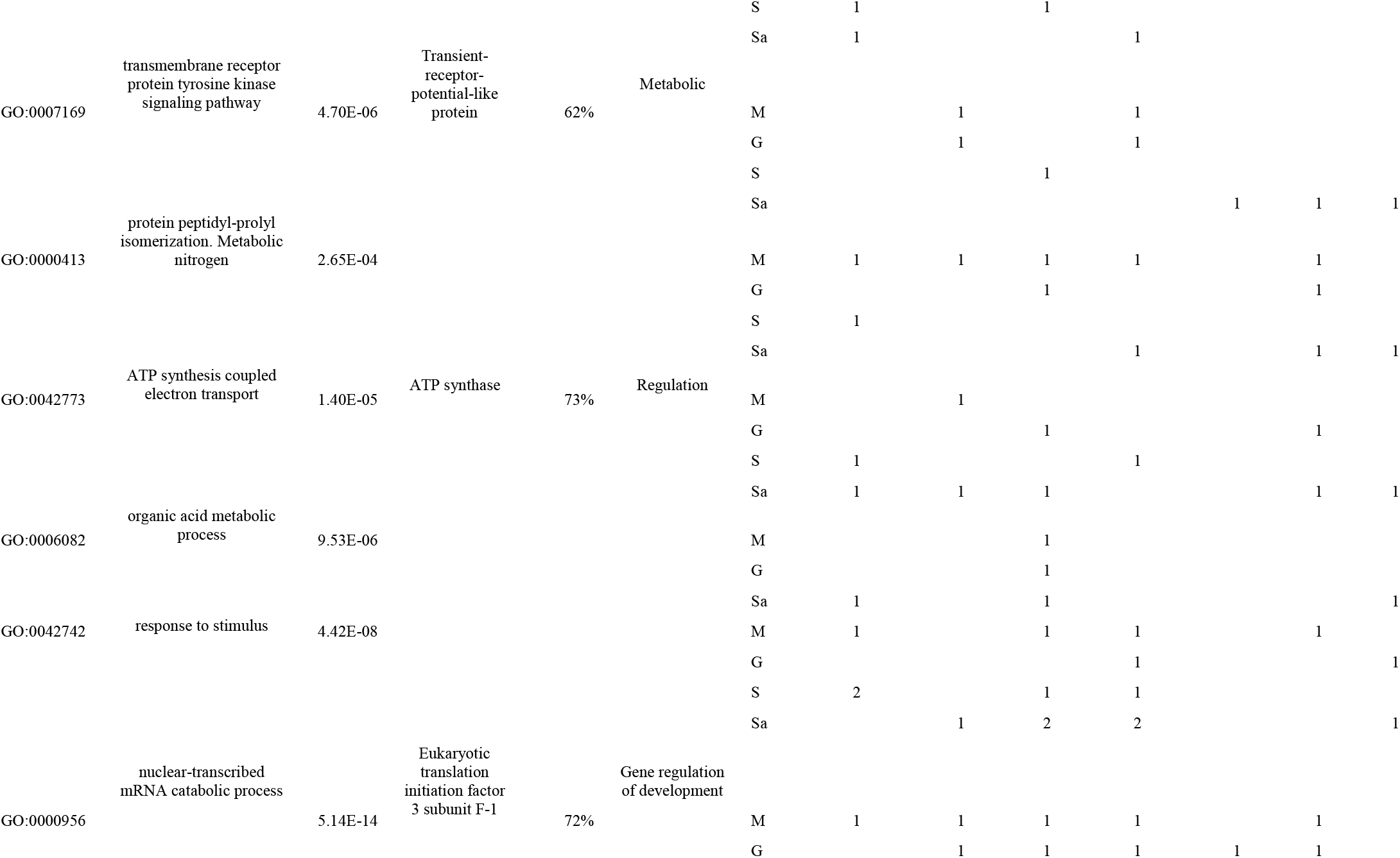
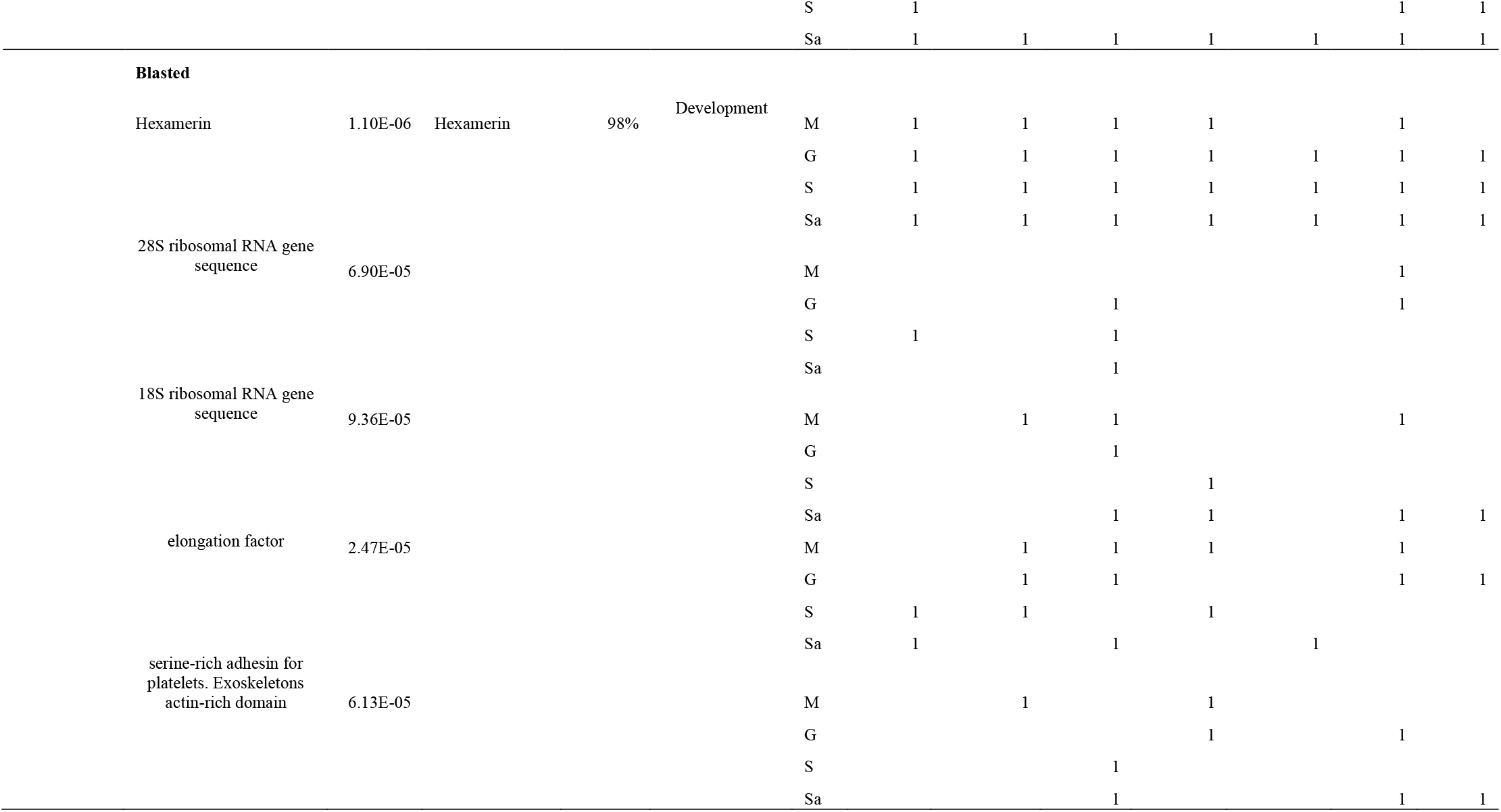
Functional annotated and blasted genes obtained from differential expression (DEs), weight gene co-expression network analysis (WGCN), and homologs species-specific analysis summarized on 30 functions and their associated protein. GO = Gene ontology, GR = geographical region, M = Matsuyama, G = Gifu, S = Sendai, Sa = Sapporo.

**Table S7.** Weight gene co-expression network analysis (WGCNA) modules with a high association to latitude across samples.

| WGCNA modules | Correlation | p-value | Associated genes |
| --- | --- | --- | --- |
| MEbrown | 0.140103 | 0.041 | 2 |
| MEred | 0.358910 | 0.007 | 4 |
| MEblack | 0.310956 | 0.012 | 5 |
| MEblue | 0.257571 | 0.020 | 2 |
| MEturquoise | 0.272889 | 0.007 | 3 |
| MEgreen | 0.272109 | 0.017 | 2 |
| MEyellow | 0.215771 | 0.009 | 3 |
| MEgrey | 0.145937 | 0.042 | 1 |

**Fig S1.**
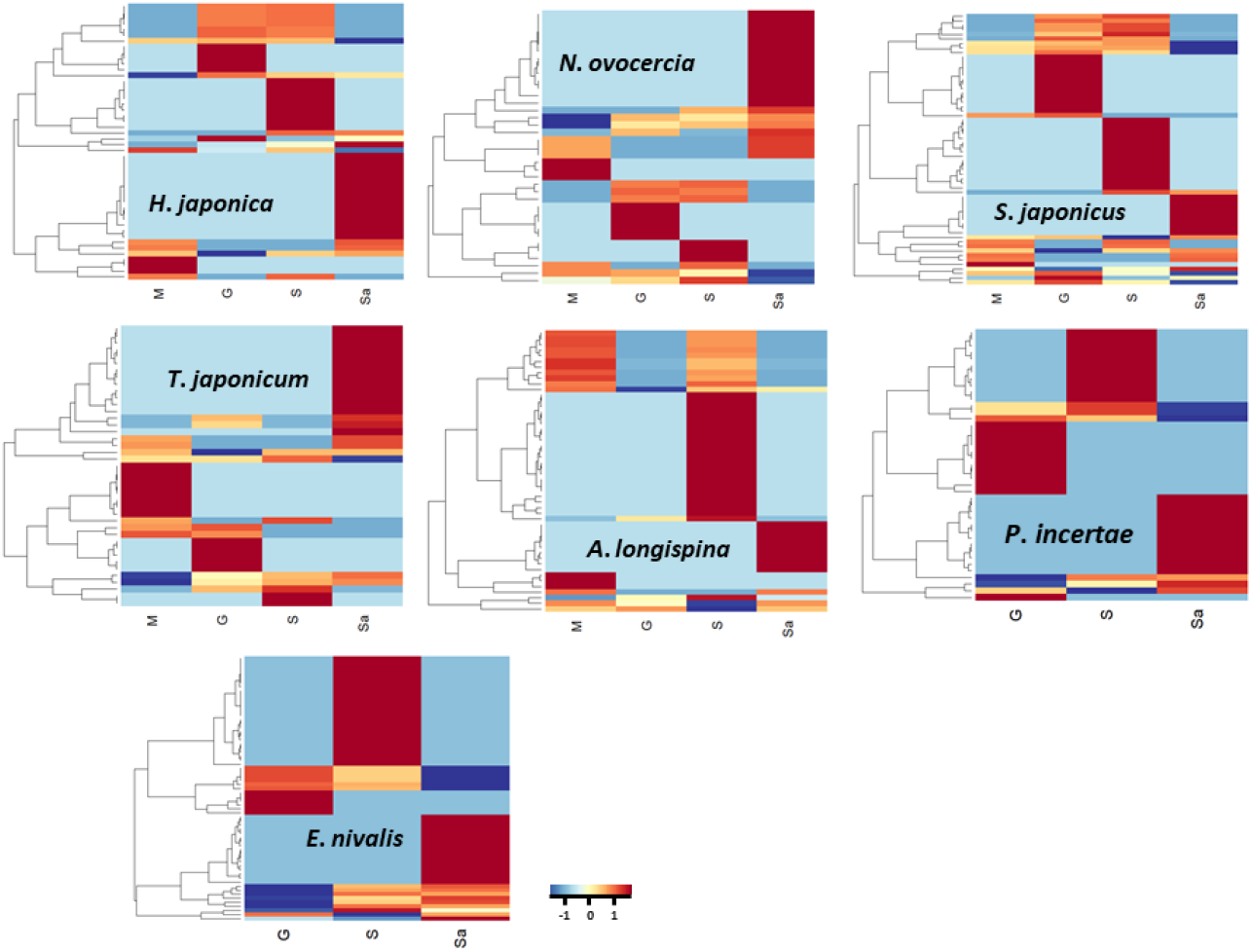
Differentially expressed genes (DEGs) of the stonefly species across 4 geographical regions in Japan. Red colors represent high expressed genes, while blue colors represent low (a color key is located at the downside). M = Matsuyama; G = Gifu; S = Sendai; Sa = Sapporo.

